# Effects of Diet on Plumage Coloration and Pigment Deposition in Red and Yellow Domestic Canaries

**DOI:** 10.1101/027532

**Authors:** Rebecca Koch, Kevin J. McGraw, Geoffrey E. Hill

## Abstract

The Atlantic Canary (*Serinus canaria*) is the most common caged bird with extensive carotenoid plumage coloration. Domestic strains of canaries have been bred for a range of colors and patterns, making them a valuable model for studies of the genetic bases for feather pigmentation. However, no detailed account has been published on the feather pigments of the various strains of this species, particularly in relation to dietary pigments available during molt. Moreover, in the twentieth century, aviculturists created a red canary by crossing Atlantic Canaries with Red Siskins (*Carduelis cucullata*). This “red-factor” canary is reputed to metabolically transform yellow dietary pigments into red ketocarotenoids, but such metabolic capacity has yet to be documented in controlled experiments. We fed molting yellow and red-factor canaries seed diets supplemented with either β-carotene, lutein/zeaxanthin, or β-cryptoxanthin/β-carotene and measured the coloration and carotenoid content of newly grown feathers. On all diets, yellow canaries grew yellow feathers and red canaries grew red feathers. Yellow canaries deposited dietary pigments and metabolically derived canary xanthophylls into feathers. Red-factor canaries deposited the same plumage carotenoids as yellow canaries, but also deposited red ketocarotenoids. Red-factor canaries deposited higher total amounts of carotenoids than yellow canaries, but otherwise there was little effect of diet treatment on feather hue or chroma. These observations indicate that canaries can use a variety of dietary precursors to produce plumage coloration and that red canaries can metabolically convert yellow dietary carotenoids into red ketocarotenoids.

---

In most birds, red, orange, and yellow plumage coloration is produced by carotenoid pigments (Goodwin 1984, McGraw 2006). No birds can synthesize carotenoids *de novo*, but some birds can biochemically modify the carotenoids that they ingest (Brush 1981). The most common carotenoids in the diets of most birds are lutein, zeaxanthin, β-carotene, and β-cryptoxanthin, and some birds convert these dietary pigments into yellow canary xanthophylls or red ketocarotenoids (carotenoids containing a ketone group) that they deposit into feathers or bare parts (McGraw 2006). Although these metabolic transformations have been deduced for a handful of birds (Brush 1990, McGraw 2006), such data are surprisingly lacking for the Atlantic Canary (*Serinus canaria*), the most common caged bird with extensive carotenoid plumage coloration.

In the wild on the Canary Islands, Atlantic Canaries have yellow feathers on their heads and breasts, with mostly brown wing, tail, and dorsal plumage (Collar et al. 2010). This appearance is retained in some domestic canary varieties, particularly those bred for singing ability. In the so-called “color-bred canaries,” however, a fantastic range of color patterns and novel shades of red, orange, and yellow have been generated through selective breeding (Walker and Avon 1993, Birkhead 2003). In our study of feather pigments, we focused on the simple lipochrome canaries, which have bright carotenoid-based coloration distributed more-or-less uniformly across the plumage with little or no melanin pigmentation of feathers (Walker and Avon 1993). Yellow lipochrome canaries have uniform bright yellow coloration across their plumage. Red-factor lipochrome canaries, which originated in the early 1900s by crossing yellow lipochrome canaries with Red Siskins (*Carduelis cucullata*; Birkhead 2003, 2014), have uniform red coloration across their plumage. Although red-factor canaries are a hybrid taxon, the hybrid offspring were back-crossed with canaries with each generation selected for red coloration but no other siskin characteristics (Birkhead 2003). Thus, individuals of this breed retain no measureable remnants of siskin phenotype except red plumage coloration.

Within the pet trade, canaries are often “color fed” by adding high doses of carotenoid pigments to the diet of molting birds to induce an especially brilliant plumage color; in fact, even yellow lipochrome canaries may acquire an orange tint upon consumption of a diet exceptionally high in red ketocarotenoids, though only red-factors may acquire truly red coloration (Birkhead 2003, Birkhead 2014). However, the diets of seed-eating songbirds like canaries typically contain exclusively yellow or orange pigments that must be converted into red pigments for ornamental coloration (Hill 2006). Isolating the red-factor canary’s ability to metabolize yellow dietary carotenoids into red carotenoids in the absence of color feeding is essential to understanding the differences in carotenoid processing between red and yellow canaries, and may have important implications generally for the control and evolution of red carotenoid metabolism in birds and other animals.

Canaries are important birds for ornithological research. They are model animals for song and neurobiological research (Goldman and Nottebohm 1983, Brenowitz and Beecher 2005), and strains have been developed with genetic knockouts for carotenoid pigmentation (white canaries) as well as for color variants (red, yellow, orange, frosted, dark red, etc.; Walker and Avon 1993). Canaries could serve as a model for studies of the genetics of carotenoid plumage coloration (Pointer et al. 2012, Walsh et al. 2012), and a first step in this process is deducing what feather pigments are produced from what dietary precursors. In the only published experimental study of feather pigmentation in domestic canaries, Brockmann and Völker (1934) detected canary xanthophylls in the feathers of canaries held on a seed diet, but did not examine the metabolic processes occurring within these birds. Modern carotenoid analytical techniques are now needed to substantiate and build on the results of this study, which was one of the first to quantify carotenoid content in the feathers of any bird.

Here, we assessed the effect of diet on plumage pigmentation in yellow and red-factor lipochrome canaries by feeding molting red and yellow canaries different carotenoid-containing diets during plumage molt and measuring the pigments deposited in newly grown feathers. Our goal was to demonstrate, under controlled experimental conditions, that red-factor canaries can convert yellow dietary carotenoids into red plumage pigments and to determine the range of dietary precursors from which canaries can produce different metabolic plumage pigments.

## METHODS

All canaries were housed as pairs in 0.46 by 0.46 by 0.92 m cages illuminated with full-spectrum lights set to the natural photoperiod of Auburn, AL. Birds were fed a maintenance diet of primarily canary seed (All Natural Canary Blend, Jones Seed Company), which provides sufficient nutrition for canaries to molt and to breed, but has low levels of yellow dietary carotenoids (<20 µg/g, lutein with low levels of zeaxanthin and β-carotene; Brockmann and Völker 1934) that are well below the supplemented levels (see more below). Birds of both sexes were included in the experiment because domestic canaries are sexually monochromatic (Koch and Hill in press).

In July 2014, we randomly divided 18 red-factor and 18 yellow color-bred lipochrome canaries (Marvin Walton, The Canary’s Nest), evenly split between both sexes, equally into three treatment groups that were supplemented with different yellow dietary carotenoids: lutein/zeaxanthin (FloraGLO, The Vitamin Shoppe), β-carotene (β-Carotene 15, GNC), or ground freeze-dried papaya (Freshlydried, Jersey City, NJ). Papaya was used as a source of β-cryptoxanthin, though it also provided birds with some β-carotene; red papaya has about four times the concentration of β-cryptoxanthin as β-carotene (Chandrika et al. 2003). We chose these diets because they provided birds with all four common carotenoids in the diets of songbirds, with each of the four dietary carotenoids supplied in abundance in at least one dietary treatment. Three birds in the lutein/zeaxanthin treatment died before the start of the experiment, so our final sample sizes were 4 red and 5 yellow in the lutein/zeaxanthin treatment, 6 red and 6 yellow in the β-carotene treatment, and 6 red and 6 yellow in the papaya treatment.

Carotenoids for all treatments were applied as powder to seed by adding about 3 g carotenoids to 20 g of canary seed. By this method of supplementation, we had no way to determine exactly how much carotenoids per day were ingested by birds. However, our methods were sufficient for the goals of this study: to determine whether red or yellow canaries have the physiological capability to convert dietary yellow carotenoids to red ketocarotenoids, and to make deductions about which dietary carotenoid are suitable precursors for carotenoid conversion.

Birds received experimental supplementation throughout the duration of their annual fall molt, ending in October 2014. We monitored birds for growing feathers, and removed 6-10 newly grown breast feathers for color measurement and carotenoid analysis. We stacked feather samples on nonreflective black cardstock paper and measured their color using an Ocean Optics USB4000 spectrophotometer and the OOIBase32 program according to standard color measurement protocols (Saks et al. 2003). From the raw reflectance spectra, we used the program CLR (v. 1.05; Montgomerie 2008) to calculate one value of hue (variable name H3) and one value of chroma (also known as saturation; S3); we selected these variables because they have been shown to most accurately reflect the carotenoid content of feathers in another cardueline finch that can display both red and yellow plumage, the House Finch (*Haemorhous mexicanus*; Butler et al. 2011).

We assessed the carotenoid content of each bird’s feather sample using high-performance lipid chromatography (HPLC) according to the methods described in Friedman et al. (2014). While our primary goal was to qualitatively describe the types of carotenoids present or absent within canaries of each treatment and color type, we also used analysis of variance (coupled with post-hoc tests using Tukey’s Honest Significant Difference method), t-tests, and linear regression models in R (version 3.0.2; R Core Team 2015) to investigate possible quantitative relationships between diet treatment, plumage color, and feather carotenoid concentrations within red and yellow canaries.

## RESULTS

Based on human visual categorization, all red-factor canaries produced only orange and red feathers (never yellow) and all yellow lipochrome canaries produced only yellow feathers, regardless of diet treatment. Yellow lipochrome canaries deposited only yellow canary xanthophylls and lutein into feathers, whereas red-factor canaries deposited yellow canary xanthophylls, lutein, and red ketocarotenoids into plumage (Table 1; Appendix). Thus, red but not yellow canaries were able to produce red ketocarotenoids from both carotenes (carotenoids containing only carbon and hydrogen; e.g. β-carotene) and hydroxycarotenoids (carotenoids containing a hydroxyl group, e.g. zeaxanthin) precursors (Fig. 1).

**TABLE 1.**
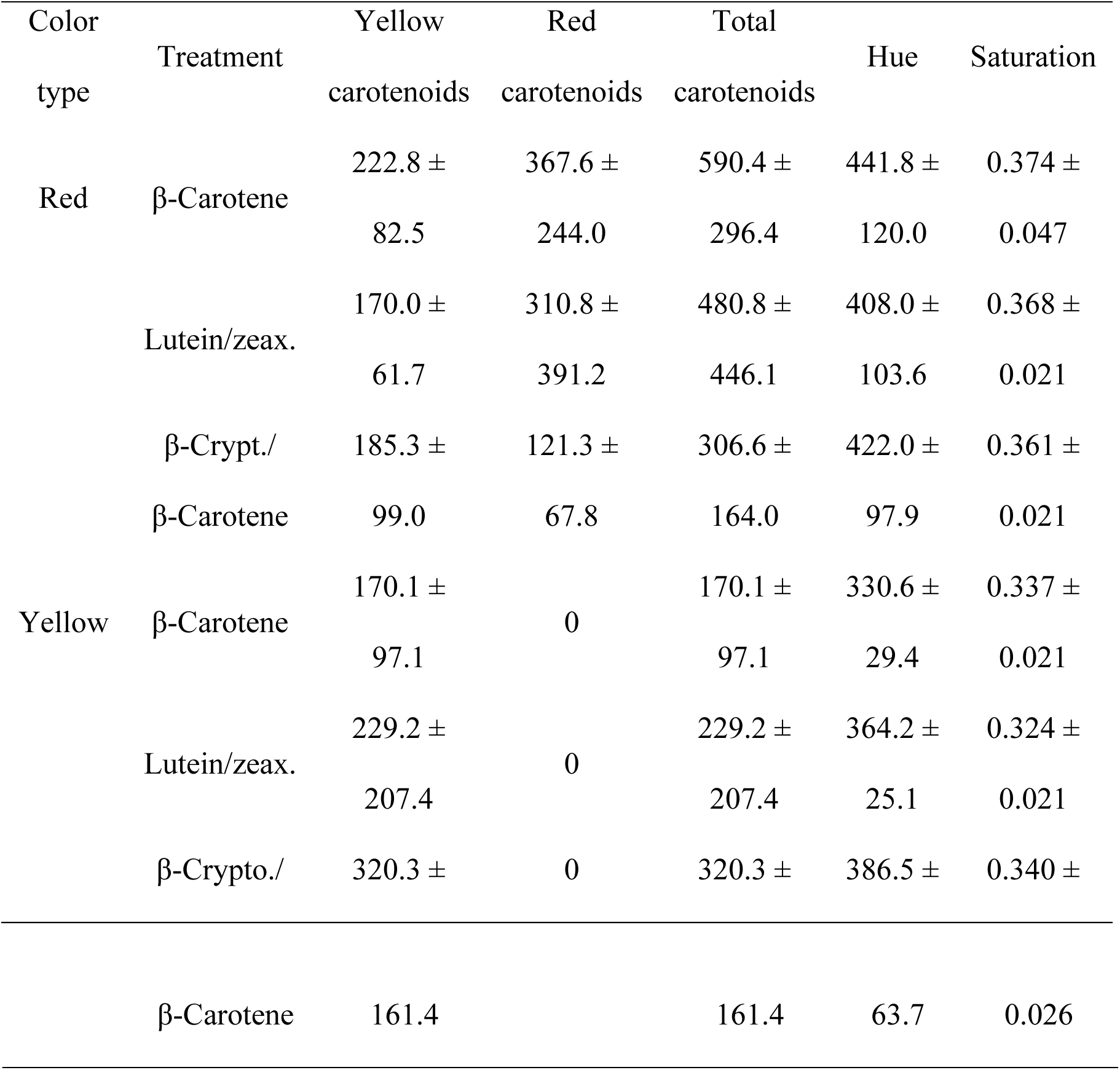
Average carotenoid content (±SD) of feathers from red or yellow canaries on one of three supplemental carotenoid treatments: β-carotene, lutein and zeaxanthin, or β-cryptoxanthin and β-carotene. All carotenoid measurements are in units of ng carotenoids per g feather; hue is in units of nm, such that higher values indicate redder color; saturation is a unitless ratio calculated from the feather reflectance spectra such that higher values indicate more saturated color. “Yellow carotenoids” comprises lutein, canary xanthophylls A, B, and C, and xanthophyll isomers generated from these pigments during extraction; “Red carotenoids” comprises echinenone, canthaxanthin, α-doradexanthin, and ketocarotenoid isomers generated during extraction.

**FIG. 1.**
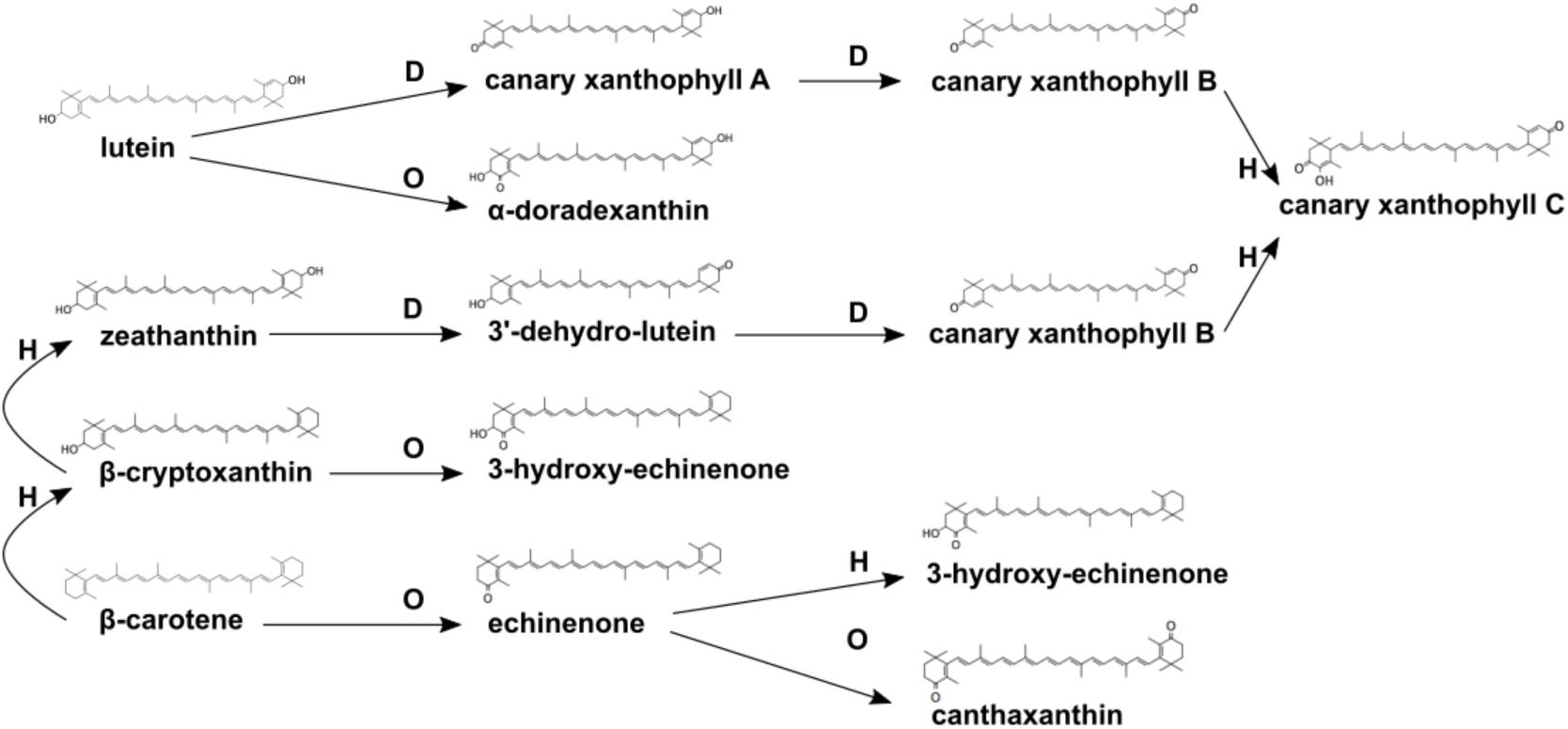
Diagram illustrating the chemical structures and relevant conversion pathways of four dietary carotenoids (lutein, zeaxanthin, β-cryptoxanthin, and β-carotene) into the metabolically altered carotenoids detected in canary feathers (canary xanthophylls A, B, and C, echinenone, canthaxanthin, and α-doradexanthin). The letters above the arrows indicate the nature of the reaction undergone: D = dehydrogenation, H = hydroxylation, and O = oxidation (McGraw 2006).

Red canaries had higher total feather carotenoid concentration than yellow canaries across all treatments (F_1,27_ = 6.84, *P* = 0.014; Table 1; Fig. 2). However, there was no significant difference in concentration of *yellow* plumage carotenoid pigments between red and yellow canaries (F_1,31_ = 0.93, *P* = 0.34; Table 1), so differences in total carotenoids appeared primarily due to the extra presence of red carotenoids in their feathers.

**FIG 2.**
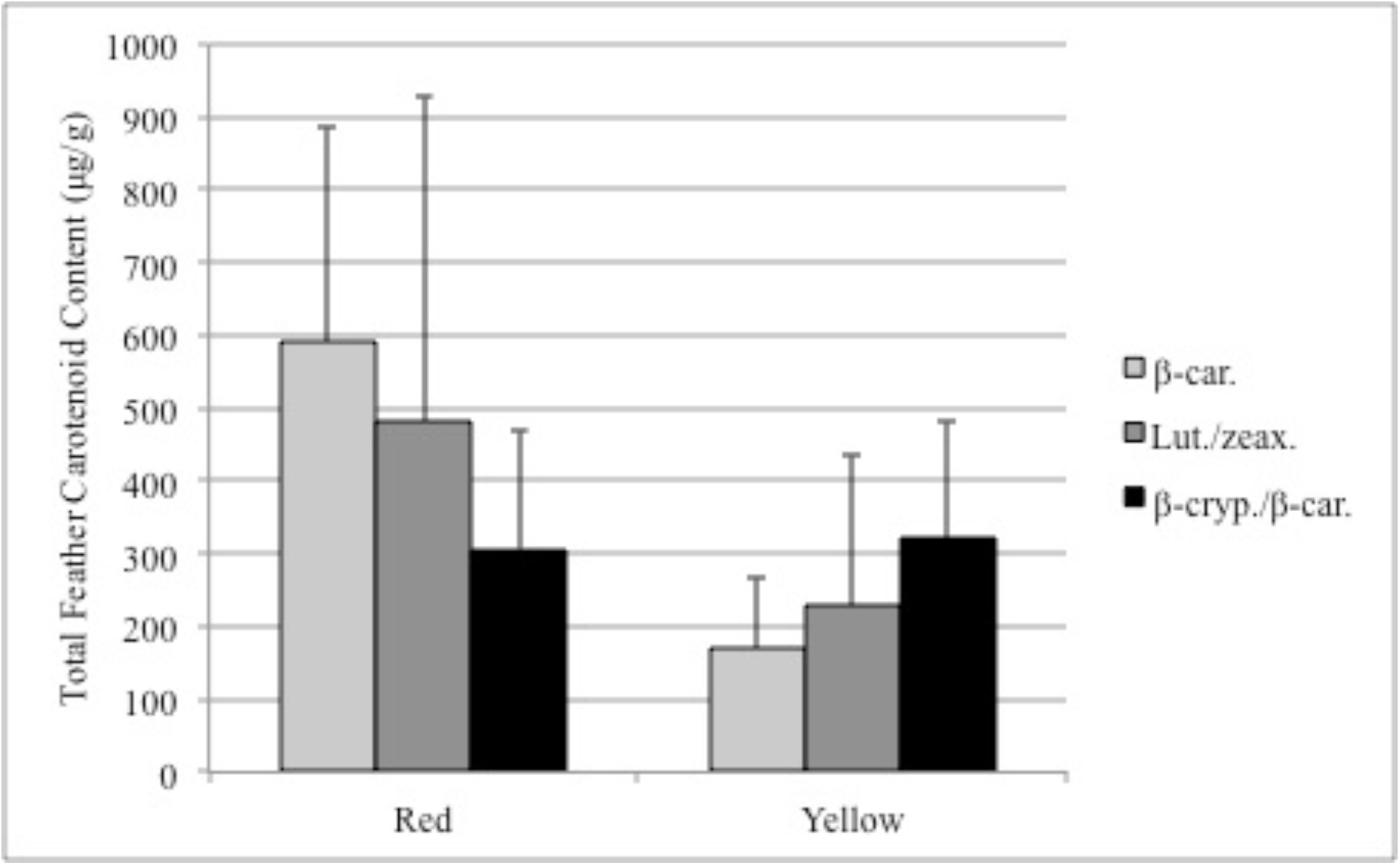
The average total feather carotenoid concentration (± SD) in the feathers of yellow and red canaries fed one of three supplemental carotenoid treatments: β-carotene, lutein and zeaxanthin, or β-cryptoxanthin and β-carotene.

There were no significant effects of diet treatment on plumage hue, saturation, total carotenoid concentration, total yellow carotenoid concentration, or total red carotenoid concentration (for red canaries only; all *P* > 0.16), with the exception that red-factor canaries on the β-carotene treatment had significantly more total carotenoids than yellow canaries on the same treatment (t = 3.30, df = 6.06, *P* = 0.016; Table 1; Fig. 2). However, we found that the total carotenoid concentration of canary feathers significantly affected plumage saturation (t = 5.30, df = 30, *P* < 0.001), but not hue (t = 1.56, df = 30, *P* = 0.13), in both color types. Saturation is therefore a better predictor than hue of feather carotenoid content in this species.

## DISCUSSION

Wild-type canaries deposit canary xanthophylls in their feathers to produce the species-typical yellow feather coloration (Stradi 1999). In the early twentieth century, aviculturists crossed Atlantic Canaries with Red Siskins to produce canaries capable of growing red feathers (Birkhead 2003, Birkhead 2014), and all domestic canaries with red plumage are descended from such siskin-canary crosses (Walker and Avon 1993). Although how red-factor canaries were genetically engineered is well known (Birkhead 2003, Birkhead 2014), the dietary bases and plumage pigment pathways of red or yellow domestic canaries have not be described (Stradi 1998, McGraw 2006). Thus, the studies presented here represent the first controlled experimental tests of the hypothesis that red lipochrome canaries produce red ketocarotenoids from yellow dietary pigments.

On any of the carotenoid diets that we provided, which varied from containing xanthophylls (e.g. lutein/zeaxanthin) to carotenes (e.g. β-carotene), yellow canaries deposited canary xanthophylls A, B, and C as well as lutein in their feathers. Thus, regardless of substrate (those four most common in the diets of terrestrial birds and ranging from yellow to orange), these birds could not metabolize dietary pigments into red plumage pigments. In addition, we found that red-factor canaries can convert a diverse array of dietary precursors—namely β-carotene, lutein/zeaxanthin, and β-cryptoxanthin—into the red ketocarotenoids α-doradexanthin, canthaxanthin, and echinenone. These data are the first experimental demonstration that red canaries can produce red ketocarotenoids from yellow dietary precursors alone.

Because we did not carefully control the quantities of carotenoids consumed by birds in our experiments, comparisons of feather carotenoid concentrations across diet treatments must be considered cautiously. Nevertheless, the data present some interesting patterns. When fed the same diets, red canaries had significantly more total carotenoids in their feathers than yellow canaries. In contrast, red and yellow canaries did not differ in yellow plumage carotenoid concentration, so the difference in total feather carotenoid levels was due to the addition of red pigments to the yellow pigments present in both canary strains. This observation implies that the pathway for ketolation of red pigments in canaries does not work from the end point of the pathway for canary xanthophylls, in which case yellow pigments would be depleted and would exist in red birds in lower concentrations. Rather, it seems that the red ketolation pathway runs in parallel to the yellow hydroxylation pathway (Hill and Johnson 2012). Parallel conversion pathways likely explain our finding that in canary plumage carotenoid concentration predicts saturation but not hue of feathers. Hue has been found to reflect the ratio of red to yellow carotenoid pigments in feathers of House Finches, which do not use parallel conversion pathways (Inouye et al. 2001); the lack of a relationship between carotenoid pigment concentration and hue in the canary indicates that the feathers contain a generally uniform ratio of red and yellow pigments in feathers, indicative of parallel conversion pathways. Saturation, on the other hand, tends to reflect the absolute concentrations of carotenoid pigments in feathers (Hill and McGraw 2006).

None of the treatments resulted in significant differences in total feather carotenoid concentrations of either yellow or red birds, except for the β-carotene treatment on which red birds had more total carotenoids than yellow bird. It is likely the lack of precision in carotenoid delivery veiled differences in how various carotenoid precursors were used to produce feather pigments, but our results do show that a range of pigment precursors can enable birds to reach the same display endpoint. This study is the first to establish that domestic red-factor, but not yellow, canaries can metabolize yellow-orange dietary pigments into red ketolated pigments.

## ACKNOWLEDGMENTS

We would like to thank R. Montgomerie for development of the CLR program. Several Auburn University undergraduates and graduate students (A. Halasz, X.R. Franko) helped with canary husbandry and supplementation. R.E. Koch was supported by NSF GRFP throughout the project.

## APPENDIX.

The concentrations in ng of all carotenoids assessed by HPLC and the hue and chroma spectrographic values of the feathers of yellow or red-factor lipochrome canaries fed one of three supplemental dietary treatments: β-carotene, lutein and zeaxanthin, or β-ccryptoxanthin and β-carotene.

